# An Automated Machine Learning Framework for Antimicrobial Resistance Prediction Through Transcriptomics

**DOI:** 10.1101/2024.06.22.600223

**Authors:** Adil Alsiyabi, Syed Ahsan Shahid, Ahmed Al-Harrasi

**Affiliations:** Natural and Medical Sciences Research Center, University of Nizwa, Birkat Al-Mouz, Nizwa, 616, Oman

**Keywords:** transcriptomics, machine learning, antibiotic resistance, AMR

## Abstract

The emergence of antimicrobial resistance (AMR) poses a global threat of growing concern to the healthcare system. To mitigate the spread of resistant pathogens, physicians must identify the susceptibility profile of every patient’s infection in order to prescribe the appropriate antibiotic. Furthermore, disease control centers need to be able to accurately track the patterns of resistance and susceptibility of pathogens to different antibiotics. To achieve this, high-throughput methods are required to accurately predict the resistance profile of a pathogenic microbe in an automated manner. In this work, a transcriptomics-based approach utilizing a machine learning framework is used to achieve this goal. The study highlights the potential of using gene expression as an indicator of resistance to different antibiotics. Results indicate the importance of starting with a high-quality training dataset containing high genetic diversity and a sufficient number of resistant samples. Furthermore, the performed analysis reveals the importance of developing new methods of feature reduction specific to transcriptomic data. Most importantly, this study serves as a proof-of-concept to the potential impact of deploying such models to reduce the mortality rate associated with AMR.

## Introduction

The overuse and misuse of antibiotics have led to a significant global threat to public health leading to antimicrobial resistance (AMR), referring to the ability of microorganisms to withstand the effects of antimicrobial drugs, rendering them ineffective [1]. The AMR presents a critical challenge that hampers the effectiveness of treatments for infectious diseases, resulting in prolonged illnesses, heightened mortality rates, and increased healthcare expenses [2]. Additionally, the rise of multi-drug-resistant pathogens has emphasized the immediate requirement for efficient strategies to confront this growing threat [3]. Given the potentially severe implications of AMR, it has become crucial to enforce rigorous measures for regulating the usage of antimicrobial agents, advocating for antimicrobial stewardship, and fostering the development of innovative therapeutic alternatives to counter the proliferation of resistance [4].

The prevalence of AMR has markedly increased on a global scale in recent years [5]. This rise underscores the persistence of the AMR challenge and emphasizes the immediate need for collaborative action to restrain its spread [6]. The collected data reveal concerning patterns, with an expanding number of pathogens demonstrating resistance to various antimicrobial agents, making the treatment of infections more complex and, at times, ineffective [7]. Projections for the upcoming decade suggest a concerning trajectory, indicating a potential surge in AMR cases if prompt preventive measures are not implemented. It is estimated that global deaths attributable to AMR could reach 10 million by 2050 if preventative measures are not put in place [8]. The persistence and amplification of AMR require immediate action through the implementation of extensive and collaborative strategies on both national and international levels [9].

The growing frequency of antimicrobial resistance demands that we shift to more personalized approaches toward the management of infections with resistant pathogens [10]. Personalized or precision medicine optimizes treatment strategies according to the needs of individual patients, that is, patient profiles, which encompass the underlying determinants of disease, including their genetic makeup, past infections, and unique specific features of the infecting organism [11, 12]. This approach is critical in the context of AMR, as it enables healthcare providers to prescribe the most effective antibiotics, thereby minimizing the risk of further resistance development [13]. The availability of new diagnostic tools, such as whole-genome sequencing and rapid phenotyping, will help identify pathogen resistance patterns faster and more accurately [14, 15]. In combination with clinical laboratory information systems, physicians can determine the level of antibiotic susceptibility and provide optimal, as well as reasonable, care [16]. Personalized treatment not only leads to better patient care but also contributes to the larger goal of reducing antibiotic use and slowing the spread of resistance. This narrow-spectrum approach to antibiotic therapy is now considered to be a cornerstone in response to AMR that involves robust diagnostic capacity and integrated health strategies to ensure that the right drug, at the right time, is targeted at the right patient [17].

Recognizing the severity and complexity of the problem, AMR surveillance has emerged as a crucial method for comprehending and tracking the prevalence and dissemination of resistance to antimicrobials [18]. While playing a critical role in tracking the dynamics of resistance patterns and guiding treatment strategies, the current AMR surveillance approaches encounter various limitations. One of the primary challenges involves the standardized methodologies and protocols for collecting, analyzing, and interpreting data [19]. This lack of uniformity impedes the accurate comparison and evaluation of AMR trends across different geographic regions and healthcare environments [20]. Furthermore, the absence of real-time monitoring capabilities hinders the timely identification of emerging resistance patterns, leading to delayed responses and the potential escalation of resistance rates [21]. Consequently, the limitations of current surveillance systems emphasize the need for new and comprehensive strategies to strengthen the monitoring and management of AMR [22].

Transcriptomics and high-throughput sequencing have emerged as leading technologies that have revolutionized the field of molecular biology, allowing for the swift and thorough examination of genetic material on an unprecedented level [23]. These advanced sequencing techniques enable the parallel sequencing of numerous DNA or RNA molecules, making it easier to investigate the genetic composition of complex microbial communities and to pinpoint genetic alterations associated with antimicrobial resistance [24]. Conversely, transcriptomics concentrates on the study of RNA transcripts within cells, offering valuable insights into gene expression patterns and molecular pathways involved in the progression and spread of resistance mechanisms [25, 26]. The integration of these advanced techniques into AMR surveillance fosters a comprehensive understanding of the genomic and transcriptomic landscapes of resistant microbes, unraveling the complex mechanisms underlying the evolution and spread of resistance [26, 27].

Leveraging the power of machine learning (ML) algorithms in conjunction with high-throughput sequencing and transcriptomic data further enhances the predictive capacity of AMR surveillance systems [28]. The upsurge in interest surrounding the integration of machine learning (ML) in healthcare has grown significantly in the last decade. This surge can be attributed to the substantial rise in the accessibility of biological and medical data, remarkable advancements in computational capabilities, and crucial breakthroughs in algorithm development [29]. By harnessing the analytical capabilities of machine learning, predictive models that identify patterns of resistance emergence [30], predict the likelihood of resistance development [28], and anticipate potential treatment outcomes, thereby enabling the implementation of proactive and targeted intervention strategies to mitigate the spread of AMR [31]. The integration of machine learning algorithms facilitates the real-time analysis of large-scale genomic and transcriptomic data, enabling the timely identification of emerging resistance patterns and the prompt implementation of tailored interventions [32, 33]. Combining the strengths of high-throughput sequencing, transcriptomics, and machine learning can foster a comprehensive and dynamic AMR surveillance framework that not only enhances our understanding of resistance mechanisms but also informs the development of effective antimicrobial stewardship programs and therapeutic interventions, ultimately contributing to the global efforts to combat the proliferation of antimicrobial resistance [34]. In this study, a machine learning-based computational approach was developed to predict the susceptibility of *E. coli* strains to different antibiotics (Figure 1). Starting with the gene expression level of only eight genes, a sampling-based automated machine-learning framework was applied to identify the optimal classification model capable of accurately predicting the susceptibility of each sample to a set of commonly prescribed antibiotics. This work demonstrates the feasibility of using gene expression to predict antibiotic resistance, a procedure that can be highly automated and which is ideal for analyzing large sample sizes. More importantly, the study highlights some important features of the transcriptomic dataset that need to be considered in future studies.

**Figure 1.**
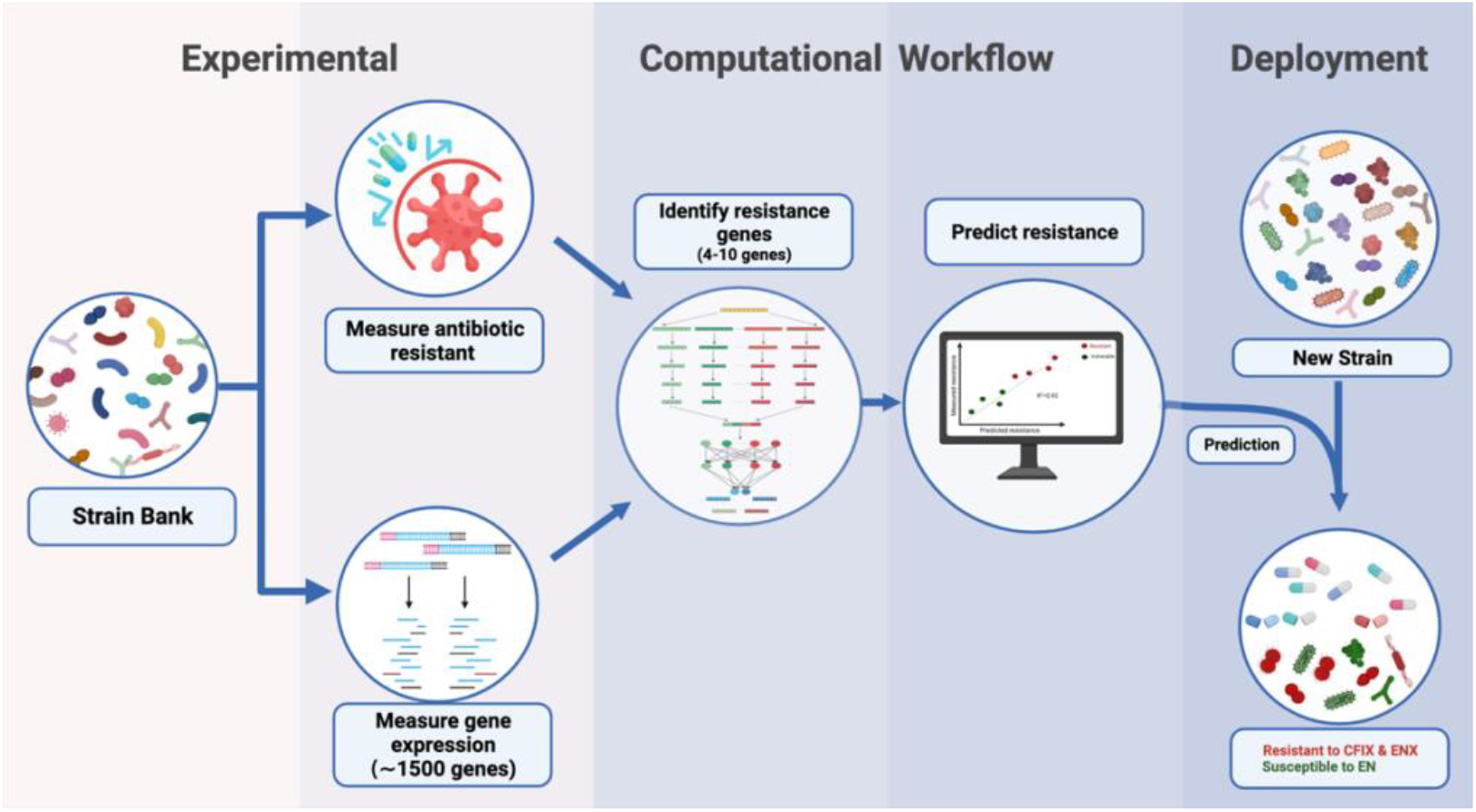
Workflow illustrating a transcriptomics-based computational approach to predict antibiotic susceptibility of pathogenic bacteria.

## Methods

### Data retrieval and preprocessing

The datasets used in this study were collected from a previously published article by Suzuki et al. [35] upon request. Namely, a dataset containing the resistance profiles of 41 strains of *E. coli* was used to create a target vector (y). The gene expression profiles were used to generate the feature matrix (X). The feature matrix and target vector were then used to train various ML models, which were subsequently used to predict the resistance of any strain given its expression profile.

In addition to the feature set containing all 4444 measured genes, three reduced feature sets were generated to determine the effects of overfitting and to test the predictive power of different subsets of genes. The first reduced feature set contained eight genes identified to be highly correlated with resistance in *E. coli* [35]. The second subset contained 64 genes determined to regulate the activity of *E. coli*’s iModulons [36]. iModulons are defined as fundamental subunits of the bacterial transcriptome, each representing a core module of bacterial physiology [37]. Finally, the third feature set comprised a combination of the first two subsets.

### Minimal inhibitory concentration (MIC) breakpoints

To convert the MIC values provided into a binary susceptible vs resistant phenotype, breakpoint values need to be chosen as a threshold [38]. Conventionally, CLSI guidelines [39] are used to identify the MIC breakpoint value for which the pathogen is considered resistant to a specific antibiotic. Supplementary Table 1 provides a summary of these values for the antibiotics considered in this study. However, the majority of measured MIC values in the utilized dataset were below the breakpoint values for the majority of antibiotics. Therefore, an arbitrary cut-off of 2 μg/ml was used to label a strain as resistant to an antibiotic.

### Model training

The sampling-based ML package auto-sklearn [40] was utilized to perform the training and model generation. Following conventional ML guidelines, 80% of the data was used to train and validate model accuracy, while 20% was held out to test model performance and generalizability [41]. Due to the limited number of samples, 3-fold cross-validation was deployed as the resampling procedure to evaluate model performance and identify the best ML algorithm [42]. During classification, data stratification was performed [43] to ensure that each fold had a similar number of resistant samples. The AutoSklearnRegressor and AutoSklearnClassifier classes were utilized to perform the regression and classification analysis, respectively. An important parameter to both classes is the training run time, which determines how long the pipeline is allowed to iterate through different ML algorithms and hyperparameter combinations. A run time of 5 hours was found to yield optimal results. During regression analysis, the optimization metric chosen to rank model performance was the F1 score, which is the harmonic mean of precision and recall [44]. Similarly, the ROC-AUC score was used during classification to ensure that the obtained models balanced between sensitivity and specificity [45]. Finally, to further optimize model performance, auto-sklearn utilizes model ensembling, where multiple generated models can be combined into an ensemble model. An ensemble size of 3 was chosen during all analyses.

## Results and Discussion

A benchmark transcriptomic dataset with corresponding resistance measurements in *E. coli* [35] was used to determine the efficacy of the proposed pipeline. We sought to accomplish a similar goal of predicting the MIC level of *E. coli* to different antibiotics based on the expression level of 8 genes, which they had identified using a genetic algorithm. Since the authors had relied on a single and rudimentary ML approach to fit the expression data, the dataset was ideal to demonstrate the importance of using a sampling approach to identify the optimal fitting method.

### An automated sampling ML approach significantly improves prediction accuracy

The majority of studies concerned with utilizing ML to analyze biological data apply a very limited number of approaches and algorithms. This limits the performance of these methods as the applied algorithms do not necessarily result in optimal accuracies. To address this shortcoming, sampling-based automated machine learning frameworks such as Auto scikit-learn has been developed [46]. These frameworks sample through a large hypothesis space in order to identify the optimal combination of data preprocessing, feature preprocessing, and model fitting methods through a Bayesian optimization approach [47]. By utilizing such a framework, we were able to significantly improve the prediction accuracies across all antibiotics compared to the original study by Suzuki *et al*., (Table 1 and Supplementary Table 2).

**Table 1.**
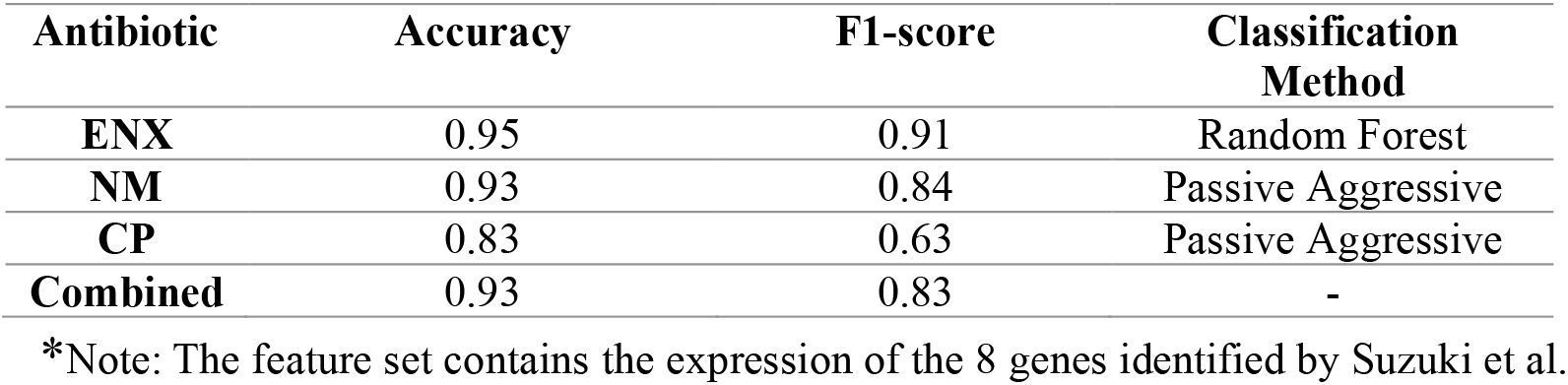
Comparison of the predictive accuracy of linear regression (LR) with automated sampling ML.

Although overall accuracy is significantly improved, model performance on unseen data remains unsatisfactory (Supplementary Table 2). This is due to two major issues. First, there is insufficient diversity in the resistance patterns of the collected samples (Figure 2). Therefore, no matter how sophisticated the learning algorithm is, it is not able to learn all patterns of gene expression required to predict the degree of resistance accurately. Second, the collected minimal inhibitory concentration (MIC) values are inherently discrete. Therefore, it is inappropriate to use a regression model to predict such values. To minimize the effect of these two issues, the resistance data was converted into a binary variable signifying susceptibility (0) and resistance (1) so that classification algorithms can be used to predict the susceptibility of each sample to different antibiotics (see methods).

**Figure 2.**
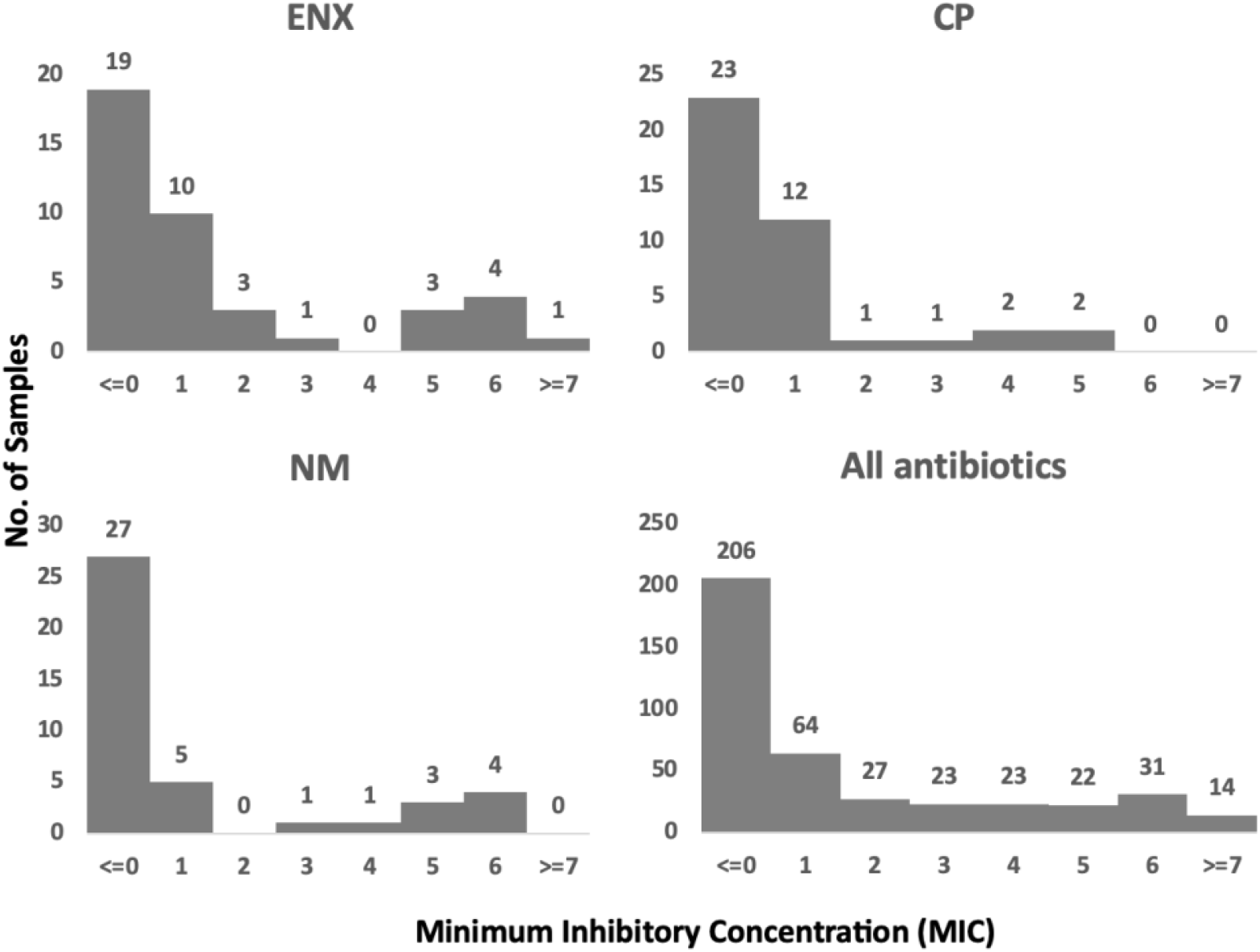
Distribution of MIC values of the *E. coli* samples used in model training.

### Accurate identification of resistant samples using minimal expression data

After converting the MIC data into resistant and susceptible samples, the classification suite of algorithms was sampled using auto-scikit learn. As can be seen from Table 2, this led to further improvements in predictive accuracy. This significant improvement is due to the simplification of the problem and the reduction of class imbalance. In the previous section, the algorithm was tasked with identifying a distinct pattern for each MIC value. However, the number of samples with high MIC values was not sufficient (Figure 1). After converting the data into susceptible vs. resistant, the algorithm had to identify one pattern only (susceptible vs. resistant). Furthermore, the samples were now divided into 2 bins only, increasing the number of samples available in each bin and therefore reducing the problem of class imbalance (Supplementary Figure 1).

**Table 2.**
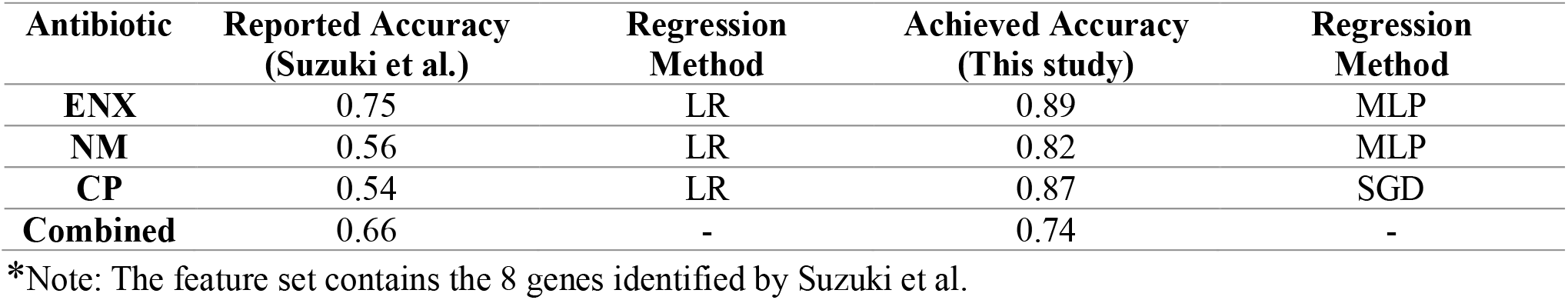
Predictive accuracy of the top-ranking classification method for different antibiotics.

In addition to improved overall accuracy, reducing class imbalance leads to better generalizability to unseen data (Supplementary Table 3). Intuitively, this can be understood by realizing that as the classes become more balanced (i.e., having a similar number of samples in each class), it becomes less likely for the model to choose the class with more samples arbitrarily [48]. Therefore, future works need to ensure a sufficient number of resistant samples in the training dataset to avoid this bias.

### The importance of identifying a reduced number of resistance genes

In addition to sampling through various fitting algorithms, automated ML frameworks can also identify the optimal dimensionality reduction approach [49]. However, dimensionality reduction does not necessarily result in a reduced number of features (i.e., genes), as many of these approaches generate combinations of existing features [50]. To determine the importance of performing feature reduction prior to applying automated ML sampling, the entire feature set of 4444 genes was used to perform classification. Interestingly, it was found that conventional methods of dimensionality reduction were not sufficient to produce models that can generalize well on unseen data for several antibiotics (Table 3 and Supplementary Table 4). This is likely due to the complexity of transcriptomic data and the high correlation in the activity of co-regulated genes. Therefore, it is essential to develop independent frameworks capable of identifying the subset of linearly independent genes that correlate with resistance. As reported [35] and demonstrated above, the use of genetic algorithms to identify such genes is promising. Furthermore, other methods, such as independent component analysis (ICA), which has been used to identify regulatory genes called iModulon regulators [36], are another promising avenue.

**Table 3.** Predictive accuracy of the top-ranking classification method using the whole transcriptome.

| Antibiotic | Accuracy | F1-score | Holdout set Accuracy | Classification Method | Feature Preprocessing Method |
| --- | --- | --- | --- | --- | --- |
| ENX | 0.90 | 0.91 | 0.56 | LDA | Nystroem Sampler |
| NM | 0.93 | 0.84 | 0.67 | SGD | PCA |
| CP | 1 | 1 | 1 | Passive Aggressive | Nystroem Sampler |
| Combined | 0.95 | 0.90 | 0.80 | - | - |

\*Note: The feature set contains the expression of all 4444 measured genes in the dataset.

### The effect of incorporating iModulon regulators to predict resistance

Although the eight genes identified by Suzuki *et al*., [35] can predict resistance with acceptable accuracy, there are other genes and pathways that independently affect antimicrobial resistance in *E. coli* but are not utilized by the model. One set of genes that are hypothesized to have a significant role to play is the set of global regulatory genes known as iModulon regulators [36].

Given the vast array of pathways regulated by these genes, it is inevitable that a subset of those pathways will have altered activity in resistant strains. Therefore, the set of 64 iModulon regulators was appended to the original eight genes, and the workflow was repeated. As can be seen from Table 4 and Supplementary Table 5, the addition of iModulon regulators improved the predictive accuracy of the holdout set (i.e., model generalizability), indicating that the expression of some global regulatory genes is indeed altered in resistant strains.

**Table 4.** Predictive accuracy of the top-ranking classification method using iModulon regulators.

| Antibiotic | Accuracy | F1-score | Holdout set Accuracy | Classification Method |
| --- | --- | --- | --- | --- |
| ENX | 0.95 | 0.91 | 0.78 | Random Forest |
| NM | 0.90 | 0.78 | 0.78 | LSVM-SVC |
| CP | 0.90 | 0.67 | 0.89 | Random Forest |
| Combined | 0.99 | 0.79 | 0.88 | - |

\*Note: The feature set contains the expression of the 64 iModulon genes and 8 genes from Suzuki et al.

It can be deduced from this analysis that feature selection is a vital component of the overall workflow. Moreover, this component of the workflow is analogous to the actively researched field of biomarker discovery [51], as both aim to identify the set of features directly related to a certain phenotype (e.g., resistance or disease). Therefore, given the rapid progress in biomarker discovery, it is expected that feature selection methods for analyzing transcriptomic data will also improve. Furthermore, as the array of biomarker discovery algorithms increases, a similar approach can be applied to automate the process of selecting resistance-causing genes. Subsequently, this can accelerate our understanding of the mechanisms underlying different types of resistance.

## Conclusion

In this study, an automated sampling-based machine learning approach was used to predict the susceptibility of *E. coli* strains to different antibiotics using the expression levels of a few genes. In addition to demonstrating the feasibility of using such an approach to identify resistant microbes in a hospital setting, the study highlights the most important aspects of the data required to obtain deployable high-efficacy models. First, the collected samples need to have a high diversity of resistant microbes so that the model can learn to identify all patterns of resistance. Second, it is vital to perform feature reduction prior to training the model, as starting with the entire set of genes leads to significant overfitting and loss in accuracy. In fact, finding generalizable feature reduction techniques for transcriptomic data is a very active area of research [52]. Furthermore, the strain bank used to generate the required transcriptomic and phenotypic data needs to contain a sufficient number of samples. Based on recent studies, the number of samples needed to perform ML on transcriptomic data is approximately ∼200 samples [35]. It is also important to ensure high genetic diversity in the strain bank to allow the model to generalize well across different regions and countries. Moreover, the type of resistance measure taken (i.e., MIC or IC_50_) can be decided based on whether the study aims to build regression or classification models. While regression models can provide predictions with higher resolution, classification models that predict susceptibility vs. resistance are often more robust and yield higher accuracies. Finally, the list of antibiotics considered needs to be determined carefully based on discussions with healthcare professionals. Figure 3 summarizes the main points to consider when designing future experiments aimed at generating datasets for machine learning analysis. Finally, while conventional approaches to identify resistance are cheaper on a per-sample basis, the potential to automate this workflow makes it very appealing for application in facilities that handle large numbers of patients.

**Figure 3.**
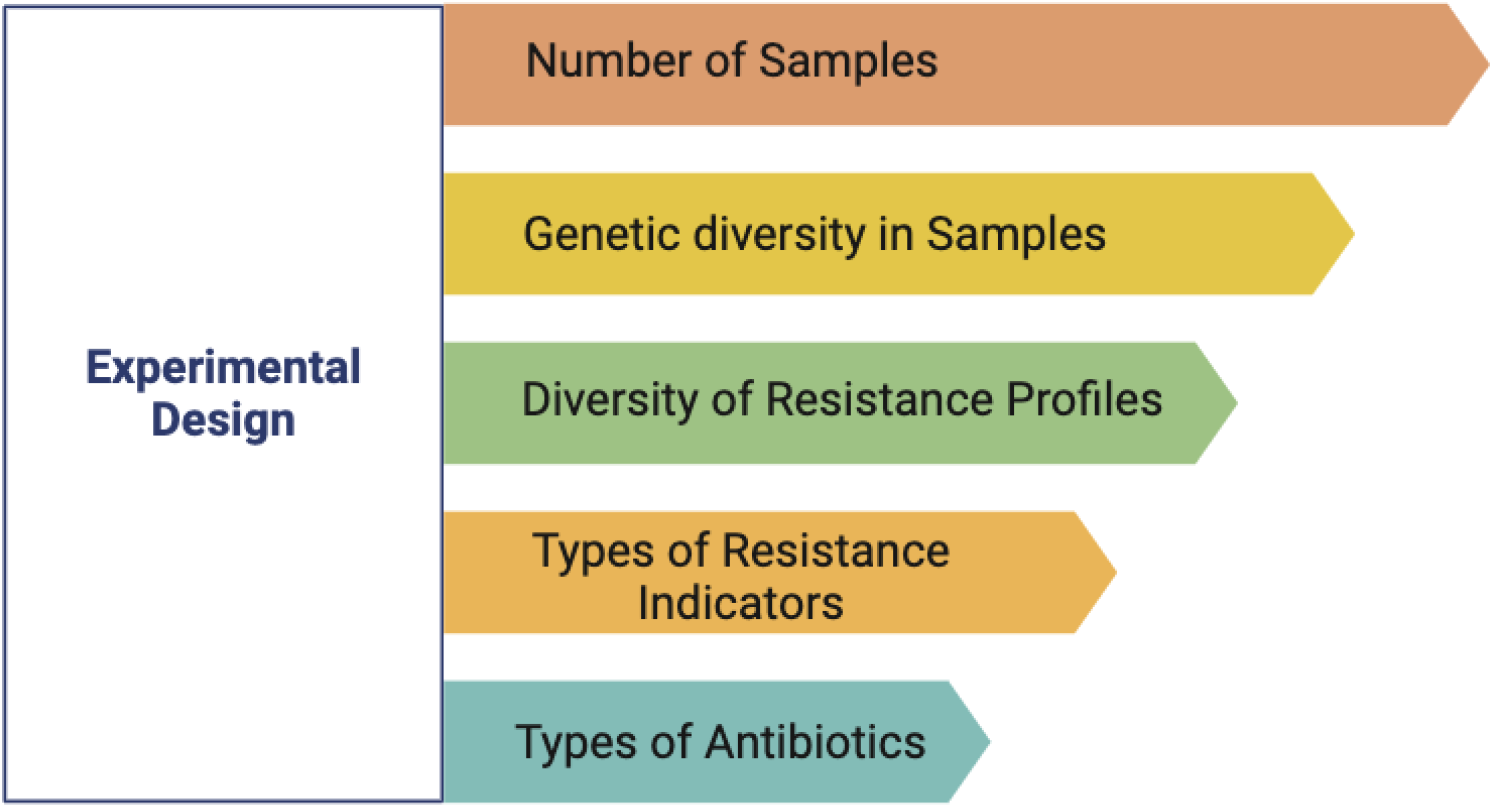
Summary of the main experimental design parameters required for robust datasets used to predict AMR.

