## Supplemental Tables and Figures for "An Automated Machine Learning Framework for Antimicrobial Resistance Prediction Through Transcriptomics"

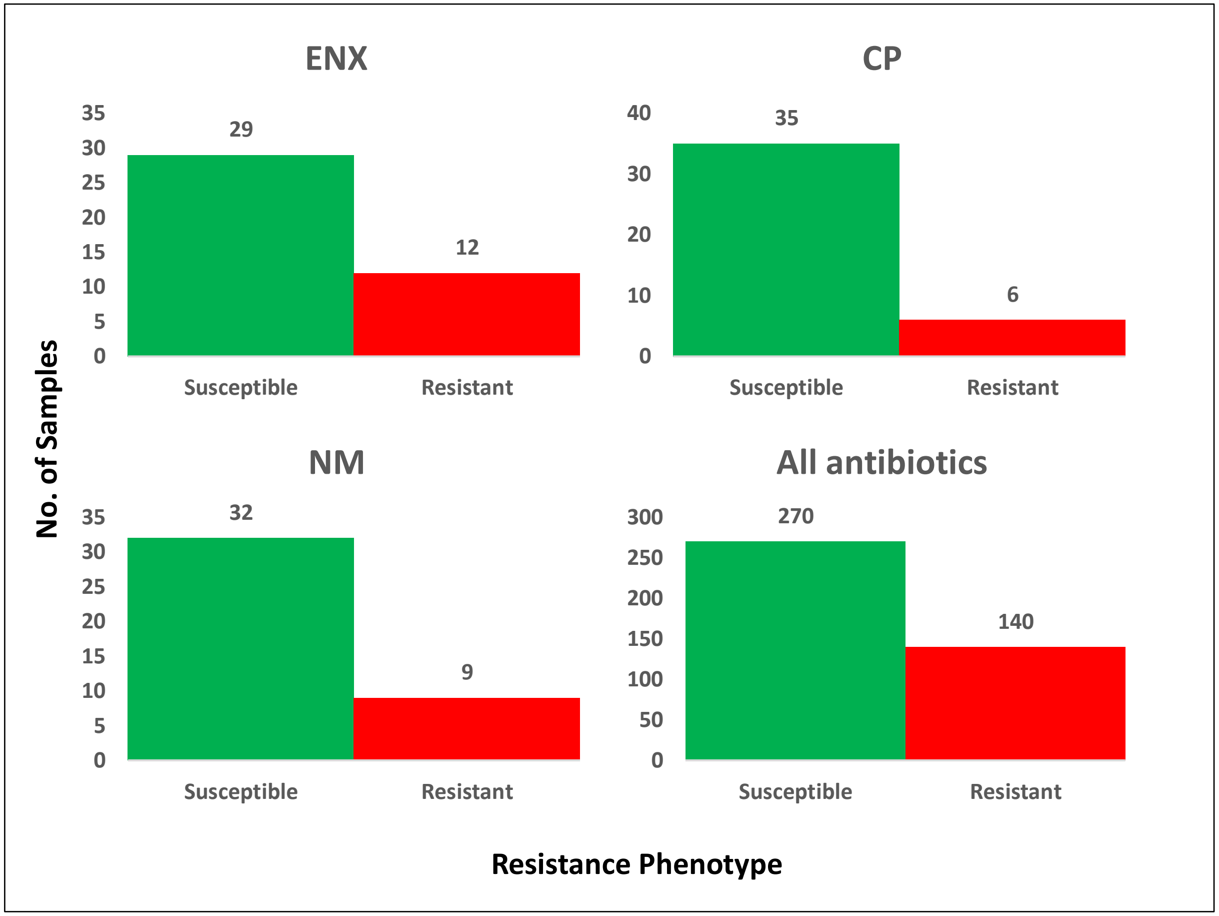


**Supplementary Figure 1. Distribution of susceptible and resistant *E. coli* samples used in model training.**

**Supplementary Table 1.** MIC breakpoint values based on CLSI guidelines

| **Antibiotic** | **Susceptible (µg/mL)** | **Intermediate (µg/mL)** | **Resistant (µg/mL)** |
|---|---|---|---|
| Cefoperazone (CPZ) | ≤16 | 32 | ≥64 |
| Cefixime (CFIX) | ≤1 | 2 | ≥4 |
| Amikacin (AMK) | ≤16 | 32 | ≥64 |
| Neomycin (NM) | - | - | - |
| Doxycycline (DOXY) | ≤4 | 8 | ≥16 |
| Chloramphenicol (CP) | ≤8 | 16 | ≥32 |
| Azithromycin (AZM) | ≤16 | - | ≥32 |
| Trimethoprim (TP) | ≤8 | - | ≥16 |
| Enoxacin (ENX) | ≤2 | 4 | ≥8 |
| Ciprofloxacin (CPFX) | ≤0.25 | 0.5 | ≥1 |

**Supplementary Table 2.** Comparison of the predictive accuracy of linear regression with automated sampling-based ML

| | Overall Accuracy | Holdout set Accuracy | Feature_preprocessor | Regressor |
|---|---|---|---|---|
| **CFIX** | 0.832757979 | 0.369959587 | feature_agglomeration | gaussian_process |
| **CPZ** | 0.937164370 | 0.665394751 | select_percentile_regression | gaussian_process |
| **CPFX** | 0.916477447 | 0.754090828 | nystroem_sampler | sgd |
| **ENX** | 0.888293583 | 0.687337546 | feature_agglomeration | mlp |
| **CP** | 0.873744575 | 0.577622555 | polynomial | sgd |
| **DOXY** | 0.397729618 | -0.32455835 | select_rates_regression | sgd |
| **TP** | 0.328678329 | 0.5677796 | extra_trees_preproc_for_regression | libsvm_svr |
| **NM** | 0.819291799 | -0.066817824 | pca | mlp |
| **AMK** | 0.639223658 | -3.62956092 | feature_agglomeration | sgd |
| **AZM** | 0.801157193 | 0.559833722 | no_preprocessing | gaussian_process |
| **Combined** | 0.74345186 | - | - | - |

**Supplementary Table 3.** Predictive accuracy of the top-ranking classifiers for each antibiotic (8 genes)

| | Overall Accuracy | F1 score | Holdout set Accuracy | Feature_preprocessor | Classifier |
|---|---|---|---|---|---|
| **CFIX** | 0.975609756 | 0.984126984 | 1 | select_percentile_classification | libsvm_svc |
| **CPZ** | 1 | 1 | 1 | extra_trees_preproc_for_classification | libsvm_svc |
| **CPFX** | 0.926829268 | 0.903225806 | 0.666666667 | nystroem_sampler | sgd |
| **ENX** | 0.951219512 | 0.909090909 | 0.777777778 | extra_trees_preproc_for_classification | random_forest |
| **CP** | 0.829268293 | 0.631578947 | 1 | select_percentile_classification | passive_aggressive |
| **DOXY** | 0.926829268 | 0.888888888 | 0.666666667 | extra_trees_preproc_for_classification | sgd |
| **TP** | 0.87804878 | 0.705882352 | 0.666666667 | feature_type | passive_aggressive |
| **NM** | 0.926829268 | 0.842105263 | 0.777777778 | feature_agglomeration | passive_aggressive |
| **AMK** | 0.951219512 | 0.857142857 | 0.888888889 | no_preprocessing | random_forest |
| **AZM** | 0.926829268 | 0.727272727 | 0.666666667 | pca | gaussian_nb |
| **Combined** | 0.92926829 | 0.84493147 | 0.81111111 | - | - |

**Supplementary Table 4.** Predictive accuracy of the top-ranking classifiers for each antibiotic (all genes).

| | Overall Accuracy | F1-score | Holdout set Accuracy | Feature_preprocessor | Classifier |
|---|---|---|---|---|---|
| **CFIX** | 0.951219512 | 0.969696969 | 0.777777778 | feature_agglomeration | extra_trees |
| **CPZ** | 0.902439024 | 0.983606557 | 1 | liblinear_svc_preprocessor | mlp |
| **CPFX** | 0.87804878 | 0.827586206 | 0.555555556 | pca | gaussian_nb |
| **ENX** | 0.902439024 | 0.909090909 | 0.555555556 | nystroem_sampler | lda |
| **CP** | 1 | 1 | 1 | nystroem_sampler | passive_aggressive |
| **DOXY** | 0.951219512 | 0.916666666 | 0.777777778 | kitchen_sinks | qda |
| **TP** | 0.87804878 | 0.705882352 | 0.777777778 | pca | sgd |
| **NM** | 0.926829268 | 0.842105263 | 0.666666667 | pca | sgd |
| **AMK** | 1 | 1 | 1 | select_percentile_classification | libsvm_svc |
| **AZM** | 0.975609756 | 0.888888888 | 0.888888889 | liblinear_svc_preprocessor | mlp |
| **Combined** | 0.93658537 | 0.90435238 | 0.8 | - | - |

**Supplementary Table 5.** Predictive accuracy of the top-ranking classifiers for each antibiotic (72 genes)

| | Overall Accuracy | F1-score | Holdout set Accuracy | Feature_preprocessor | Classifier |
|---|---|---|---|---|---|
| **CFIX** | 0.87804878 | 0.91525423728 | 0.888888889 | feature_agglomeration | mlp |
| **CPZ** | 1 | 1 | 1 | extra_trees_preproc_for_classification | random_forest |
| **CPFX** | 0.975609756 | 0.96969696969 | 0.888888889 | select_rates_classification | libsvm_svc |
| **ENX** | 0.951219512 | 0.90909090909 | 0.777777778 | feature_agglomeration | random_forest |
| **CP** | 0.902439024 | 0.66666666666 | 0.888888889 | extra_trees_preproc_for_classification | random_forest |
| **DOXY** | 0.951219512 | 0.92857142857 | 0.777777778 | polynomial | passive_aggressive |
| **TP** | 0.902439024 | 0.66666666666 | 0.888888889 | feature_agglomeration | libsvm_svc |
| **NM** | 0.902439024 | 0.77777777777 | 0.777777778 | liblinear_svc_preprocessor | lda |
| **AMK** | 1 | 1 | 1 | extra_trees_preproc_for_classification | lda |
| **AZM** | 0.951219512 | 0.74999999999 | 0.888888889 | kitchen_sinks | lda |
| **Combined** | 0.94146341 | 0.85837247 | 0.87777778 | - | - |

*Note: The feature set contains the 8 genes identified by Suzuki et al. and the 64 iModulon genes.
